# TP53-META, a meta-analysis tool for comparative transcriptomics of TP53 dependency: Examples from target silencing and liver fibrosis

**DOI:** 10.1101/2025.10.14.682117

**Authors:** Ayse G. Keskus, Eren Kumak, Merve Vural-Ozdeniz, Abdul Moiz Aftab, Said Tirkayi, Thomas Darde, Ozlen Konu

## Abstract

**Background:** TP53 is the most frequently mutated transcription factor (TF) in sporadic cancers; and its targets exhibit dysregulation at the level of expression in both cancer and non-cancer pathologies. However, there is not yet a web-based tool that enables meta-analysis and visualization of TP53-related gene expression datasets, although several databases exist to access and annotate TP53 target information. To address this gap, we developed TP53-META, an interactive R Shiny-based web tool that allows users to upload and simultaneously analyze user’s or integrated public RNA-seq datasets for effects of TP53 depletion and/or induction on the transcriptome.

**Results:** TP53-META can be used to visualize significant expression clusters as well as TF-TF, pathway-pathway, disease-gene and treatment-gene networks to determine TP53 dependency of selected treatment contrasts. We demonstrated the utility of TP53-META through two case studies. In the first, using an in-house RNA-seq data from MCF7 cells treated with siRNAs against CHRNA5 and TP53, we identified TP53-independent and dependent transcriptomic changes by CHRNA5 depletion by comparing with selected public datasets in TP53-META. In the second, we demonstrated the user data upload functionality of TP53-META before meta-analysis and extracted commonly modulated TP53-related genes in liver fibrosis using public RNA-seq datasets. TP53-META is available at http://konulabapps.bilkent.edu.tr:3838/TP53-Meta1.5/

**Conclusions:** By facilitating meta-analysis, clustering, and network-based visualizations, TP53-META enables researchers to efficiently integrate and explore TP53-related transcriptomic datasets from diverse sources, and help uncover robust expression patterns, and investigate context-specific TP53 functions.

## BACKGROUND

TP53, which is frequently altered and extensively studied in different sporadic cancers, plays an important role in the transcription of genes involved in DNA repair, senescence, and cell metabolism while acting as the “guardian of the genome” (1, 2). However, there is not yet a statistical analysis and visualization tool which can help decipher whether a treatment/process depends on TP53’s actions on the transcriptome. Moreover, TP53’s extensive contribution in multiple processes necessitates integration of multiple resources in an online web server/database.

There are already a few existing tools that compile TP53 targets or cistromes, e.g., Targetgenereg.org (3) and the P53-Cistrome shiny application (4). Despite their utility, these tools are not without limitations. For example, Targetgenereg.org lacks a repository and is not interactive. While both Targetgenereg.org and the P53-Cistrome offer TP53 specificity and meta-analysis, neither includes methods for differential gene expression (DGE) analysis. Furthermore, none of these tools can perform enrichment analysis of functional terms, transcription factor (TF) target enrichment, or network visualization in combination.

Meta-analysis is a powerful statistical methodology that allows integration and comparison of findings from different sources (5, 6) and hence can lead to the identification of context-dependent TP53 actions. Accordingly, a TP53 data-specific online tool that integrates and enables meta-analysis of TP53-related (e.g., testing of TP53 siRNA and activity inducers) RNA-seq datasets could be highly useful to researchers in the field. Moreover, due to TP53 target gene lists differing significantly from one another given the type of tissue, cell, or analysis method (7), the extraction of consistent patterns of expression would help identify biomarkers.

Meta-analysis of RNA-seq data is currently possible with different web tools. For example, DRPPM-EASY (8) allows integration of more than one dataset for comparisons via logFC comparison plots. Network Analyst/ExpressAnalyst, Omics View, and Omics Playground are highly comprehensive tools with the ability to integrate and visualize multiple datasets (9, 10, 11, 12). However, no existing tool allows comparison of multiple contrasts from one or more datasets with a filtering capability within an individual dataset filtering and biological theme comparison. In addition, visualization of treatment-gene, pathway-pathway, or TF co-membership networks by themselves or all together is also needed. Most importantly, none of the available tools are specific to modulations of TP53 activity. Although existing tools can be used to analyze one or more TP53-relevant datasets they do not contain built-in TP53-modulated datasets and TP53-related gene lists. Thus, while generic tools exist, they do not capture the unique biology, variability, or curated datasets for TP53.

To address the lack of a comprehensive and modular RNA-seq analysis tool offering TP53 specificity, we developed TP53-META. This novel tool allows DGE analysis, functional analysis, and prioritized visualization of gene-treatment interactions across four contrasts simultaneously using in-house, default public, and/or user uploaded datasets. Recently, interest in TP53’s role in breast cancer prognosis has increased due to the prevalence of TP53’s or its regulators’ mutations or dysregulation (13, 14). Moreover, multiple new players in breast cancer have been identified with interactions with TP53 based on gene silencing and knockout studies (15, 16). Accordingly, in the present study, we used our novel in-house RNA-seq dataset (GSE301555) and strongly demonstrated that the functional outcomes of CHRNA5 depletion were partly TP53-dependent. In addition, we showed in another case study that liver fibrosis stage was associated with a common set of TP53 targets based on meta-analysis of three different public datasets (GSE135251, GSE162694, and GSE174478).

## IMPLEMENTATION

TP53-META is an online application that allows the analysis of user uploaded raw RNA-seq counts and/or integrated datasets. It is mainly divided into three pages: 1) “Home” page that summarizes the integrated datasets; 2) “Upload” page that enables integrating user datasets; and 3) “Analysis” page that enables filtering, comparisons, and downstream visualizations (Figure 1). The “Analysis” page is further divided into “Data Summary” and “Results” subsections. The former allows the examination of the selected datasets by customizable volcano, bar/box, and PCA plots. The latter provides nine different sub-tabs, i.e., Analysis, Scatter Plot, Heatmap, Functional Profile, DoRothEA, Treatment-Gene Networks, MsigDB, Compare Cluster, and Correlation.

**Fig 1.**
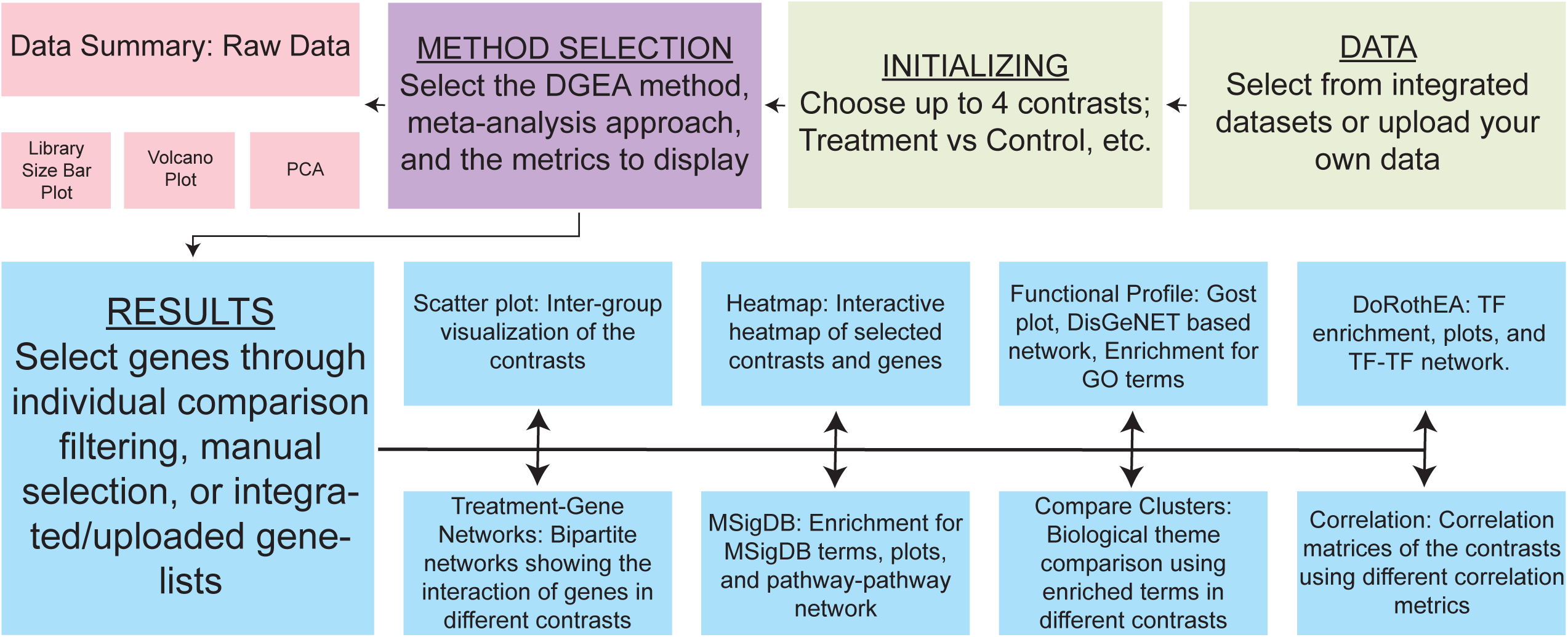
Simplified workflow of the TP53-META.

Each analysis performed by TP53-META offers customization with the selection of user-defined thresholds on analysis statistics, e.g., p-values, log fold changes, as well as customizable visualizations. Explanations on how to use each of these functions can also be found on the online tutorial page of TP53-META. The details of each tab are further explained below.

### Input Files and Datasets

TP53-META includes 13 datasets, all of which are presented on the “Home” page of the application (Table 1). Briefly, our in-house RNA-seq experiment is unique to the TP53-META application and consists of three separate treatments of MCF7 cells, i.e., an siRNA for the CHRNA5 or TP53, or a combination of these two siRNAs (GSE301555). While the application allows analysis of differences between treatments, a baseline siRNA control group is also present. The remaining datasets have been obtained from Gene Expression Omnibus (17). These include three different MCF7 cell line studies with Nutlin treatments (GSE47042, GSE86219, GSE248670) and two other datasets using MDA-MB-231 and MCF10A, each treated with an siRNA for TP53 (GSE68248, GSE74493). TP53-META also includes four gastrointestinal cancer experiments performed on different cell lines with TP53 mutations or siRNAs against a specific target and/or TP53, or treatment with Nutlin (GSE173364, GSE59841). In addition, three other datasets on other cell types focusing on TP53 activation or target silencing in the presence or absence of TP53 (GSE83635, GSE84877, GSE124508). Such a dataset collection makes it possible to test the degree of TP53 dependency of a given treatment, such as gene silencing.

**Table 1.**
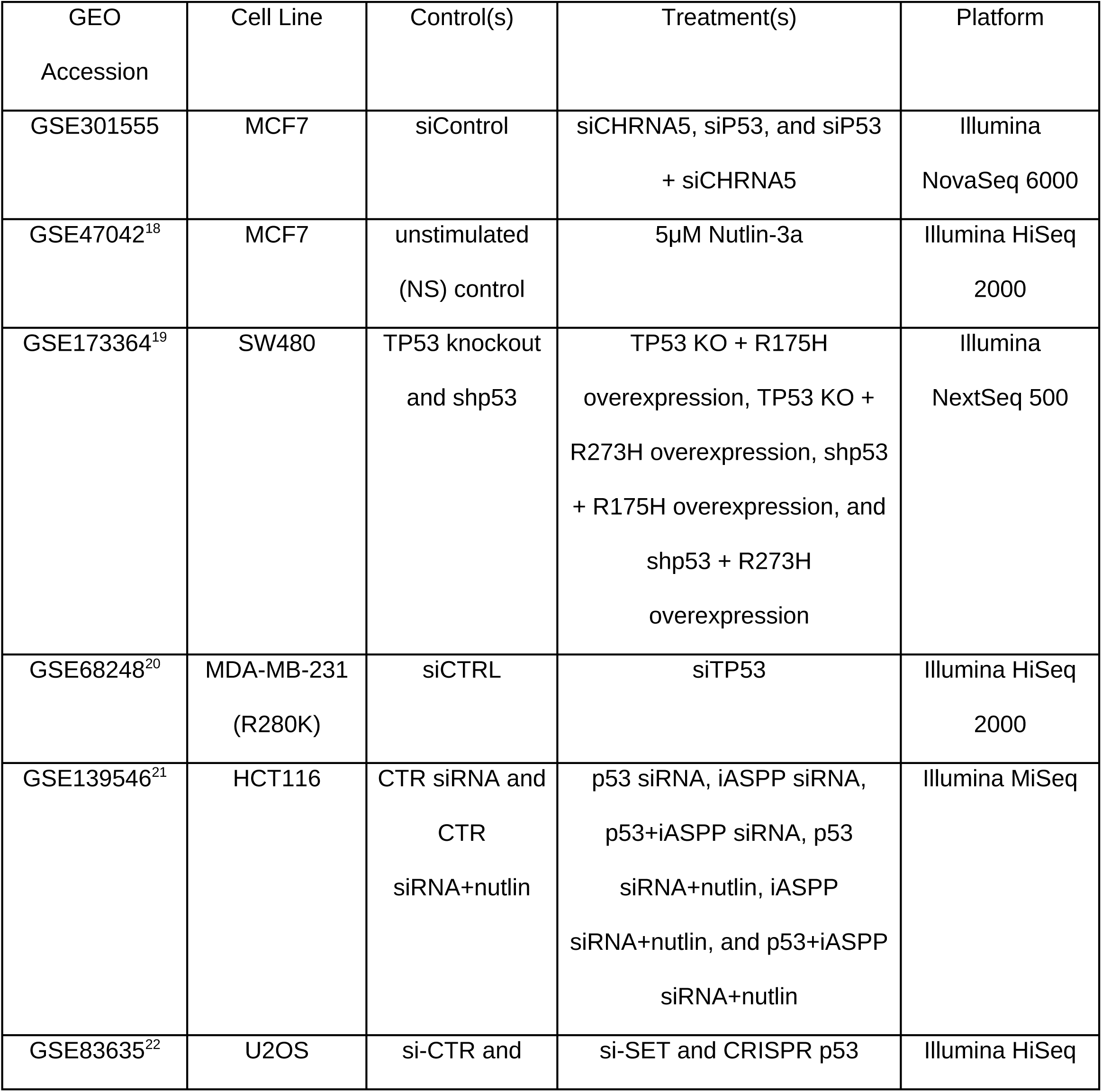
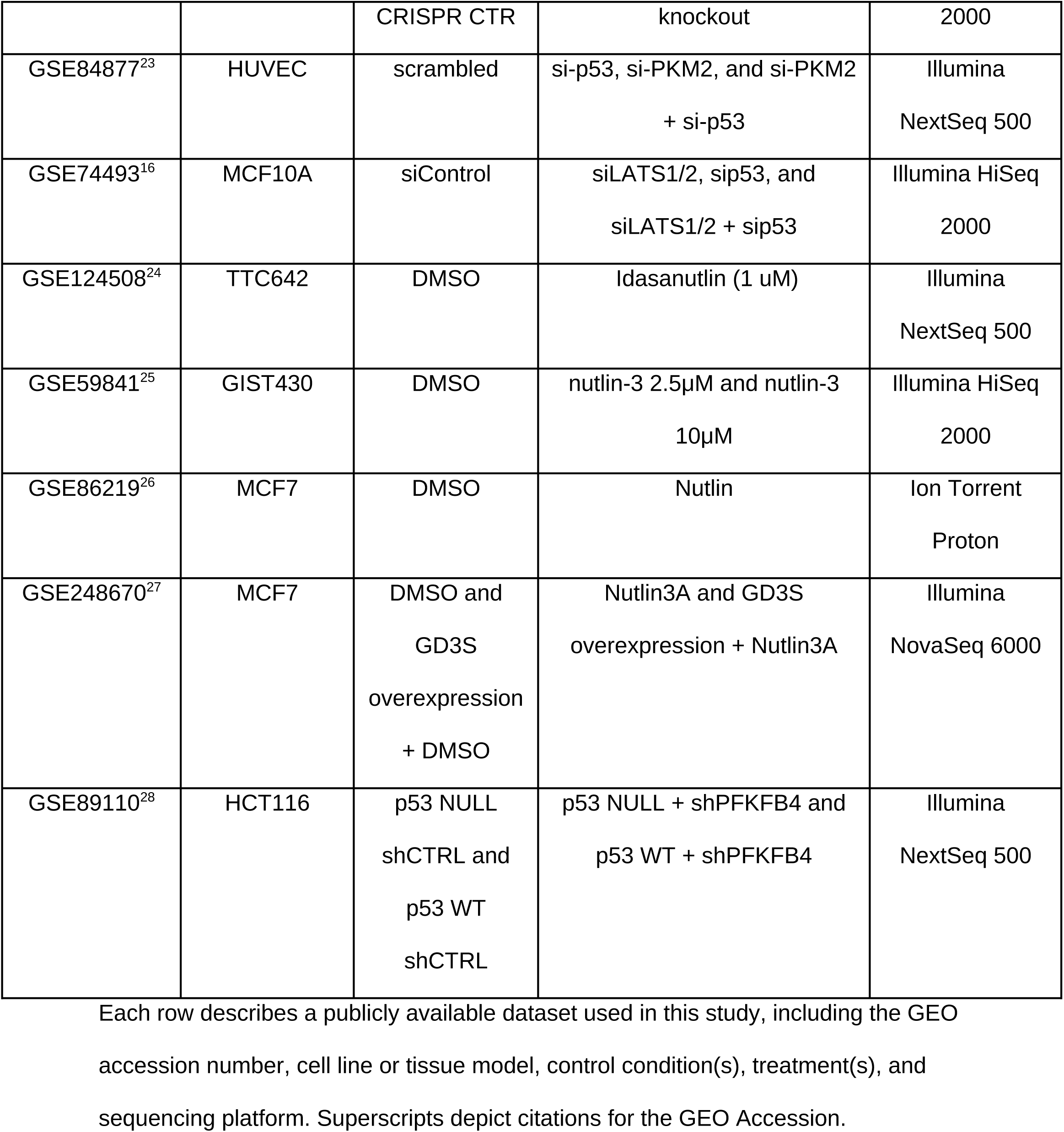
Summary of included datasets in TP53-META.

| GEO<br>Accession | Cell Line | Control(s) | Treatment(s) | Platform |
| --- | --- | --- | --- | --- |
| GSE301555 | MCF7 | siControl | siCHRNA5, siP53, and siP53<br>+ siCHRNA5 | Illumina<br>NovaSeq 6000 |
| GSE47042<sup>18</sup> | MCF7 | unstimulated<br>(NS) control | 5μM Nutlin-3a | Illumina HiSeq<br>2000 |
| GSE173364<sup>19</sup> | SW480 | TP53 knockout<br>and shp53 | TP53 KO + R175H<br>overexpression, TP53 KO +<br>R273H overexpression, shp53<br>+ R175H overexpression, and<br>shp53 + R273H<br>overexpression | Illumina<br>NextSeq 500 |
| GSE68248<sup>20</sup> | MDA-MB-231<br>(R280K) | siCTRL | siTP53 | Illumina HiSeq<br>2000 |
| GSE139546<sup>21</sup> | HCT116 | CTR siRNA and<br>CTR<br>siRNA+nutlin | p53 siRNA, iASPP siRNA,<br>p53+iASPP siRNA, p53<br>siRNA+nutlin, iASPP<br>siRNA+nutlin, and p53+iASPP<br>siRNA+nutlin | Illumina MiSeq |
| GSE83635<sup>22</sup> | U2OS | si-CTR and | si-SET and CRISPR p53 | Illumina HiSeq |
|  |  | CRISPR CTR | knockout | 2000 |
| GSE84877<sup>23</sup> | HUVEC | scrambled | si-p53, si-PKM2, and si-PKM2<br>+ si-p53 | Illumina<br>NextSeq 500 |
| GSE74493<sup>16</sup> | MCF10A | siControl | siLATS1/2, sip53, and<br>siLATS1/2 + sip53 | Illumina HiSeq<br>2000 |
| GSE124508<sup>24</sup> | TTC642 | DMSO | Idasanutlin (1 uM) | Illumina<br>NextSeq 500 |
| GSE59841<sup>25</sup> | GIST430 | DMSO | nutlin-3 2.5µM and nutlin-3<br>10µM | Illumina HiSeq<br>2000 |
| GSE86219<sup>26</sup> | MCF7 | DMSO | Nutlin | Ion Torrent<br>Proton |
| GSE248670<sup>27</sup> | MCF7 | DMSO and<br>GD3S<br>overexpression<br>+ DMSO | Nutlin3A and GD3S<br>overexpression + Nutlin3A | Illumina<br>NovaSeq 6000 |
| GSE89110<sup>28</sup> | HCT116 | p53 NULL<br>shCTRL and<br>p53 WT<br>shCTRL | p53 NULL + shPFKFB4 and<br>p53 WT + shPFKFB4 | Illumina<br>NextSeq 500 |

Each row describes a publicly available dataset used in this study, including the GEO accession number, cell line or tissue model, control condition(s), treatment(s), and sequencing platform. Superscripts depict citations for the GEO Accession.

The web framework also allows users to upload their own RNA-seq counts. This can be done using the “Upload” page, where two files are required: the raw counts and metadata including a list of conditions. The raw counts file is a CSV file with gene symbols included as the first column and raw count values of each sample in the following columns. A condition file is a one-column TXT file without a header, including the group names in the counts file in the same order. Example files for upload are also available in downloadable format, combined with interactive tables visible on the “Upload” page. Upon successfully uploading the documents, RNA-seq data as well as options for comparisons become available under the “Your Data” tab on the “Analysis” page of the application.

### Differential Gene Expression and Meta-analysis

TP53-META allows simultaneous differential gene expression of up to four contrast/comparison(s). This is done on the data selection page upon clicking on the “Analysis” button, where the application contains four dropdown menus to select the experiment, followed by the selection of the control and the treatment. The user can select limma or DESeq2 as their method of differential gene expression analysis (29, 30). Gene symbols are stored as Ensembl stable IDs using org.Hs.eg.db package and BiomaRt (31, 32). Statistics from the resulting analysis, such as log fold change (LogFC), average expression, p-value, adjusted p-value, t-value, and B-value, are displayed in the table given the user’s choice. When more than one contrast is present for the analysis, TP53-META calculates meta-p-values in addition to the p-values of individual experiments based on the poolr package (33) which provides “Fisher’s” as the default but other options can be selected, e.g., “Stouffer’s,” “Inverse chi-square,” “Binomial test,” “Bonferroni,” and “Tippett’s.” The default parameters of each method as implemented in poolr are used, with the function receiving only the vector of p-values obtained from DGEA as input.

### Data Summary

When the analysis is complete, the “Data Summary” section becomes available on the “Analysis” page, which includes raw statistics for each contrast selected, separately. The first subsection of this part contains a volcano plot of the selected contrast, where the user can manipulate the visualization by adjusting the value cutoff, logFC cutoff, point size, and label size. The second subsection is the “Plots” option, which includes library size for the data as a bar plot and principal component analysis (PCA) plot of the data colored according to the conditions. Finally, “Raw counts” are available in the last part of the “Data Summary” section and can also be downloaded for use outside of the web application.

### Filtering

The “Results” section upon completing a differential gene expression process with TP53-META contains a general “Analysis” sub-tab. A table containing all the desired statistics per gene according to previously made user selections along with meta-p-values is visible on this tab. TP53-META further allows users to display and filter only selected genes found in integrated gene lists from a dropdown menu, titled “Display Gene List”, or upload their own gene list to analyze only a selected number of genes. Integrated gene lists include TP53 related gene sets, i.e., cell cycle genes, DREAM targets, MMB-FOXM1 targets, lin37 dependent TP53 targets, lin37 independent TP53 targets, RB E2F targets, TP53 associated E2 response genes, and TP53 independent E2 response genes, which are obtained from previously conducted research (34, 35, 36). In addition to directly uploading or selecting a gene list, TP53-META offers three different ways to filter genes: 1) clicking on the row of gene of interest; 2) using the value entry options in each column to set thresholds; or 3) using the buttons “Select Upregulated” and “Select Downregulated” to extract genes, where user-defined p-value and logFC cutoff thresholds are applied to selected contrasts. When the user filters genes to use in downstream analyses, a second table also appears under the “Analysis” sub-tab, containing only the selected genes. Both tables visible under the “Analysis” tab are downloadable using the “Download” buttons below the tables. There are also “Copy” buttons to copy tables to the clipboard and a search bar to specifically search for a gene.

### Scatter Plots

The “Scatter Plot” sub-tab of the TP53-META enables users to create scatter plots of selected statistics, which can be used for comparisons within or between contrasts. While the scatter plots created by this method are interactive, they can also be customized by adjusting the point sizes. While such plots can be created for both selected genes or the entire dataset, linear regression is also applied to the data. The formula of the best-fitting line and the R^2^ value are available on the plot. TP53-META performs regression using R’s built-in linear model fitting function.

### Heatmaps

TP53-META creates interactive heatmaps using the heatmaply package so that the created heatmaps can be zoomed in for specific sections, and values are displayed upon hovering over cells (37). Created heatmaps, include row and column dendrograms with customizable distance and clustering methods. User can select from multiple methods of distance and linkage, as specified in the package. Besides these method options, the user can also select the number of clusters using a sliding bar. Heatmaps created in TP53-META are fully downloadable as HTML or PNG due to the built-in properties of the heatmaply package.

### Functional Profile

TP53-META “Functional Profile” sub-tab under the “Analysis” tab enables term enrichment for selected genes. After opening this tab, two plots can be generated: 1) a GOSt plot for the GSEA results (38); and 2) a gene-disease association network/ heatmap using the DisGeNET database (39). While creating a gene-disease association network and heatmap, TP53-META offers parameter customization where the user can change minGSSize, p-value cutoff, q-value cutoff, number of terms to display, category size scaling factor, and whether to use circular layout. The results of the GO terms enrichment analysis are presented in table format, where the user can change the minimum number of genes for testing (40). This table can be copied, printed, and downloaded as a CSV or XLSX file with the buttons above the table. The user can also visualize the enrichments using a dot plot or an enrichment map/network, or a Cnet plot of genes and terms, with colored gene nodes (41). Layout for these visualizations can be selected from multiple available methods listed as options while “circle” is the default. Finally, the number of displayed genes can be customized using the sliding bar.

### Transcription factor - Transcription Factor (TF-TF) Networks of DoRothEA

TP53-META integrates the DoRothEA database to analyze interactions between transcription factors and target genes using the DoRothEA R package (version 1.20.0) (42). If the user selects genes in the “Analysis” sub-tab, the DoRothEA output is directly calculated when the tab is opened. The DoRothEA gene sets are further fed into the hypeR package in R to calculate the significance of enrichment (43). All of these results are available in a table, ranked by their significance. This sub-tab further creates a dot plot that displays each TF’s FDR value. In this plot, sizes of dots reflect the relative number of genes targeted by the given TF.

Other functions of the “DoRothEA” sub-tab become available when one or more TFs are selected by clicking on rows of the table. When TFs are selected, a heatmap that shows expression logFC values of TF targets across the selected contrasts becomes available. If more than one TF is selected, the heatmap also includes row annotations showing the DoRothEA confidence level (A–E or NA) for each TF–target interaction. These confidence levels, obtained from the DoRothEA database, indicate the strength of literature or experimental evidence supporting the regulatory link (A = highest confidence, E = lowest). The confidence level is displayed alongside each target in the heatmap as a separate annotation column. This heatmap can be further customized by adding dendrograms for samples and genes, where the distance method can be selected. Other customization options include column text angle, row font size, and column font size.

A network of TF-TF connected based on their overlap in content (i.e., gene number) can be created upon filtering according to the maximum number of TFs or selected FDR threshold values. This network uses the Jaccard distance to represent dissimilarity between overlapping targets of the two TFs, with a user-adjustable threshold to trim the network.

In the resulting network, edges represent the number of shared genes between TFs. Hovering over an edge displays the number of overlapping genes and the associated Jaccard distance. Edge color intensity reflects the number of shared genes, with darker edges indicating more overlap. Node coloring is based on the FDR value of each TF— darker nodes indicate smaller FDRs. Node size can be adjusted based on either the FDR value or the number of overlapping genes, depending on user preference.

Additional controls include the ability to stop network motion and to display only connected nodes after thresholding. Font size is also user-adjustable. The network can be further highlighted by clicking on individual nodes or selecting them by transcription factor ID.

### Treatment-Gene Networks

The “Treatment-Gene Networks” sub-tab of the TP53-META application visualizes nodes of selected genes from the “Analysis” sub-tab alongside the treatment contrasts as a bipartite network. The networks are generated using the visNetwork R package (44) in which the gene’s expression logFC value is used for annotating the edge width when connected to a specific treatment, while edge color indicates direction of regulation, red for upregulated and blue for downregulated genes.

In TP53-META, genes and treatment contrasts are represented as separate nodes where gene node color intensity increases with the number of significant contrasts associated with that gene. TP53-META can generate such networks with up to 600 genes, a limit set to reduce crowding and network generation time. Network layout is user-selectable, with three options available: Force-based, Hierarchical - Vertical, and Hierarchical - Horizontal. Users can filter the network to include only upregulated or downregulated genes and can apply a threshold based on absolute logFC to trim the network. It is also possible to trim edges that are not significant (P ≥ 0.05). Font size is user-adjustable; and users can control network motion and toggle the inclusion of unconnected nodes using sidebar checkboxes. Selected nodes can be highlighted either by clicking on them or by choosing them through the “Select by ID” option.

### MsigDB Enrichment

Using the “MSigDB” sub-tab of the application, the user can select from different gene sets in the database (45), e.g., H (hallmark gene sets). When the analysis is complete using the msigdbr R package, TP53-META shows the results in a table format. Upon user selection, genes of selected terms can be further visualized by creating a heatmap using the logFC values of the associated genes in each contrast, and additional coloring to indicate in which geneset the gene is included. While the heatmap is interactive, the user is also able to customize it by deciding whether to include dendrograms or not, and the method of distance.

A network of the resulting enriched terms can be generated based on an FDR cutoff defined by the user. Depending on the user’s selection, the network can include only the pathways that meet the threshold or also include associated pathways that share genes with the filtered set. In these networks, pathways are represented as nodes, and edges indicate the number of shared genes between any two pathways. Node color reflects significance, with red nodes and edges representing pathways with q-values < 0.05, and blue representing non-significant ones. Node size is scaled according to the pathway’s p-value, which is also displayed when hovering over a node. Like other networks in TP53-META, this network is interactive and customizable, allowing users to select nodes, adjust font sizes, control network motion, and choose from different pathway collections.

### Compare Clusters

The “Compare Clusters” sub-tab of TP53-META allows comparison of GO enrichment results for upregulated and downregulated genes in all contrasts set by the user for comparison. A dot plot is created after this sub-tab runs the GO enrichment analysis for upregulated and downregulated genes in all contrasts using the selected genes from the “Analysis” sub-tab. In this plot, the dot sizes correspond to the GeneRatio column of the analysis, and colors correspond to p-values. While this dot plot enables users to compare enrichment results of different contrasts, it also offers customization by changing font size, absolute logFC threshold, and category number fed into the plotting function.

### Correlation Plots

The final sub-tab of the TP53-META application is the “Correlation” sub-tab. This tab utilizes the built-in correlation function of R to display the correlation coefficient between contrasts using the logFC values of selected genes in the “Analysis” sub-tab. The results of the correlation are fed into the corrplot package in R to create circular correlation plots with correlation coefficients (46). The correlation method, which includes “Pearson,” “Kendall,” and “Spearman,” is up to the user to select using the drop-down menu on the page. While the user can directly use the selected genes, it is also possible to filter for genes that are significant in all contrasts using the checkbox on the page.

### RNA-seq for siRNA-treated MCF7 cells and qPCR validations

MCF7 cells (2 × 10⁵), seeded in 6-well plates, were treated after 24h with the following for 72h: (i) control siRNA (20 nM; FlexiTube, SI03650318, Qiagen), (ii) siRNA targeting CHRNA5 (10 nM; FlexiTube, SI03051111, Qiagen), (iii) siRNA targeting TP53 (10 nM; FlexiTube, SI02623747, Qiagen), or (iv) a combination of CHRNA5 and TP53 siRNAs (each at 10 nM) using HiPerFect transfection reagent (6:1000, reagent: total volume; 301704, Qiagen). To ensure consistent siRNA concentration across groups, single siRNA treatments were supplemented with 10 nM control siRNA. Total RNA extraction (RNeasy Mini Kit, 74104, Qiagen) and cDNA generation from 1 μg of RNA (RevertAid First Strand cDNA Synthesis Kit, K1622, Fermentas) were followed by RNA sequencing on the NovaSeq 6000 platform, resulting in 150 bp paired-end reads.

Raw FASTQ files were uploaded to the Seven Bridges Cancer Genomics Cloud (47) for analysis. Upon quality control, performed using FastQC (48), reads were aligned to the human genome (hg19) using the STAR aligner with default parameters (49), and HTSeq-count was used for quantification at the gene level for Ensembl Gene IDs and genomic coordinates (50). Validation of expression level changes of selected genes was performed by quantitative reverse transcription PCR (RT-qPCR) (SYBR Green Master Mix, Roche, 04707516001) on a LightCycler 480 system (Roche) using TPT1 as the reference gene.

### Western Blot Analysis

Proteins were harvested using a freshly prepared lysis buffer composed of NaCl, Tris- HCl, NP-40, 10% SDS, protease inhibitors, and PhosSTOP (Roche), followed by protein quantification (Thermo Scientific, Cat. No. 23227). Lysates, mixed with 4× sample buffer (Bio-Rad, 161–0747) and heated for 5 minutes, were electrophoresed on 10% polyacrylamide gels (Bio-Rad, 161–0183) before being transferred onto PVDF membranes (Roche, 3010040) using a semi-dry blotting system. Membranes, blocked with 5% bovine serum albumin in TBS containing 0.2% Tween-20 (TBST), were incubated overnight at 4°C with primary antibodies against CHRNA5 (Santa Cruz, sc-376979), TP53 (Santa Cruz, sc-126), and GAPDH (Santa Cruz, sc-47724); washed and treated with HRP-linked secondary antibodies (anti-mouse: Cell Signaling, 7076; anti-rabbit: Cell Signaling, 7474) for 1 hour at room temperature. Signal detection was performed using chemiluminescence reagents and visualized with the Amersham™ Imager 600.

## RESULTS

### Applications of TP53-META to integrated or uploaded public datasets

We have used two case studies to demonstrate the use of TP53-META in different contexts. The first case study involves the use of our integrated in-house custom RNA-seq data on singular and combinatorial treatments of siRNAs against CHRNA5 and TP53 genes in MCF7 cells and their meta-analyses with integrated public datasets. The second case study shows how to upload a set of public datasets and then extract TP53 signatures that are common to all or a specific contrast.

### Case Study 1: Effects of CHRNA5 depletion on the TP53 target expression in breast cancer

CHRNA5 is a ligand-gated ion channel and a nicotinic acetylcholine receptor (nAChR) gene (51). While the role of CHRNA5 in lung cancer progression and the resistance to chemotherapeutic agents was shown before, we recently demonstrated that CHRNA5 depletion by siRNA resulted in transcriptomics changes similar to those induced by the TP53 inducer Nutlin (15). However, this was not confirmed by an application of TP53 siRNA together with a CHRNA5 siRNA before. Hence, in a new study, we performed an RNA-seq experiment using four groups: i) control siRNA; ii) CHRNA5 siRNA; iii) TP53 siRNA; and iv) combination of siRNAs [GSE301555; see Additional File 1]. Use of TP53-META enabled us to establish a significant association between CHRNA5 depletion and TP53 signaling by performing meta-analysis against integrated datasets focusing on TP53 depletion or activation by Nutlin.

### Meta-analysis of expression profiles of siTP53-treated cell lines identifies a core TP53 depletion signature

First, expression profiles of our siTP53 treated MCF7 cells (GSE301555) were compared against those of two publicly available siTP53 datasets obtained from cancer cell lines carrying WT TP53, i.e., MCF10A breast cancer cell line (GSE74493), HCT116 human colon cancer cell line (GSE139546). In addition, a dataset using MCF7 (GSE47042) cells, treated with Nutlin; a commonly used MDM2 inhibitor for activation of TP53, was also included as a control (Figure 2A). Genes significant in all datasets upon differential gene expression using Limma-Voom (p-value < 0.05 in each study, as well as meta-p-value obtained by Fisher’s method < 0.05) revealed that while all siTP53 treatment comparisons showed positive pairwise correlations, the Nutlin-treated MCF7 cells showed a negative correlation across all siTP53-treated experimental groups, as expected. (Figure 2B).

**Fig 2.**
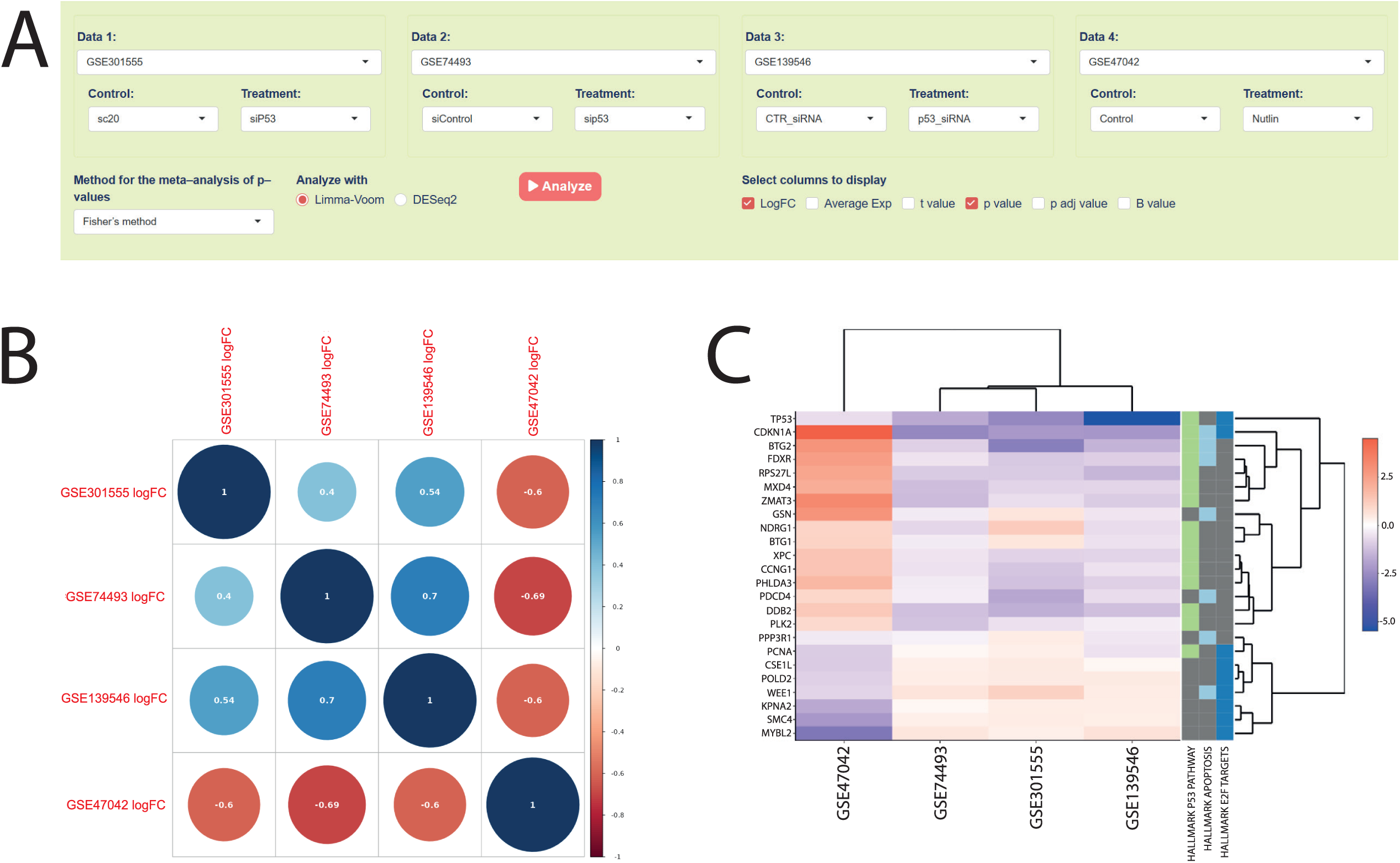
**A)** Selected comparisons and configurations for siTP53 analysis across different datasets in TP53-META. **B)** Pearson correlation matrix based on log fold change (logFC) values of 94 genes commonly altered in all four datasets used. **C)** Heatmap for pathway analysis results on selected MSigDB Hallmark pathways.

A total of 94 genes were identified when genes significant in all comparisons were considered [see Additional File 2A]. According to the TF analysis based on DoRothEA, genes were enriched in transcription factors TP53 and E2F4 [see Additional File 2B].

TP53 target genes in DoRothEA exhibited negative logFC values in siTP53 experiments and positive logFC values in the Nutlin treatment, except for *BTG1*, *NDRG1*, and *PCNA*. *BTG1* and *NDRG1* had positive logFC values in siTP53 (GSE301555) and Nutlin-treated MCF7 cells, whereas *PCNA* was downregulated in siTP53-treated HCT116 cells and Nutlin treatment. Both transcription factors and gene relationships were classified as score A in the DoRothEA classification indicating their confidence levels were high.

Furthermore, the MSigDB analysis showed that filtered genes were enriched in the hallmark TP53 pathway, apoptosis, and E2F targets (Figure 2C).

Moreover, to show the gene list-specific use of TP53-META, the same analysis was performed by uploading and filtering according to the TP53 census target gene list obtained from Fischer et al. (52). For these genes (n = 270, meta-p-value using Fisher’s method < 0.05), the functional profile for DisGeNET using logFC values of our in-house data included TP53 downregulation-related diseases such as xeroderma pigmentosum, skin carcinogenesis, and endometrial adenocarcinoma [see Additional File 3]. GO terms enrichment analysis also resulted in apoptotic, DNA damage, TP53 regulation, and cell cycle-related processes [see Additional File 4]. Accordingly, TP53-META has made an effective analysis of different studies simultaneously while allowing visualization and downstream analysis of significant genes obtained from the differential gene expression analysis.

### Depletion of CHRNA5 in MCF7 cells mimics the expression signature of Nutlin treatment

Based on our previous findings (15, 53), we used TP53-META to investigate whether CHRNA5 depletion by siCHRNA5 treatment exhibited similarity to the treatment with Nutlin (Figure 3A). All public Nutlin treatment studies included in the application, i.e., GSE47042, GSE248670, and GSE86219, were performed in the MCF7 breast cancer cell line, as our siCHRNA5 experiment from GSE301555. These datasets were reanalyzed using Limma-Voom, followed by Fisher’s method to perform a differential gene expression analysis with meta-p-values. We then filtered across all the datasets based on the user-uploaded TP53 census gene list, filtered for significance (p < 0.05 for all comparisons). Accordingly, we obtained a gene set with 94 genes, different from the set obtained in the abovementioned siTP53 comparisons. Within these 94 genes, only eight showed logFC values in different directions across datasets. These included *PTPRE*, *RNF144B*, *KRT15*, *ANXA4*, *HES1*, and *COBLL1*, showing negative logFC values in siCHRNA5 treatment, as well as *TRIP6* and *GPC1* having mixed directions across the analysis. The “Scatter Plot” and “Correlation” tabs of the application showed that siCHRNA5 treatment had a positive correlation with Nutlin treatment [see Additional File 5].

**Fig 3.**
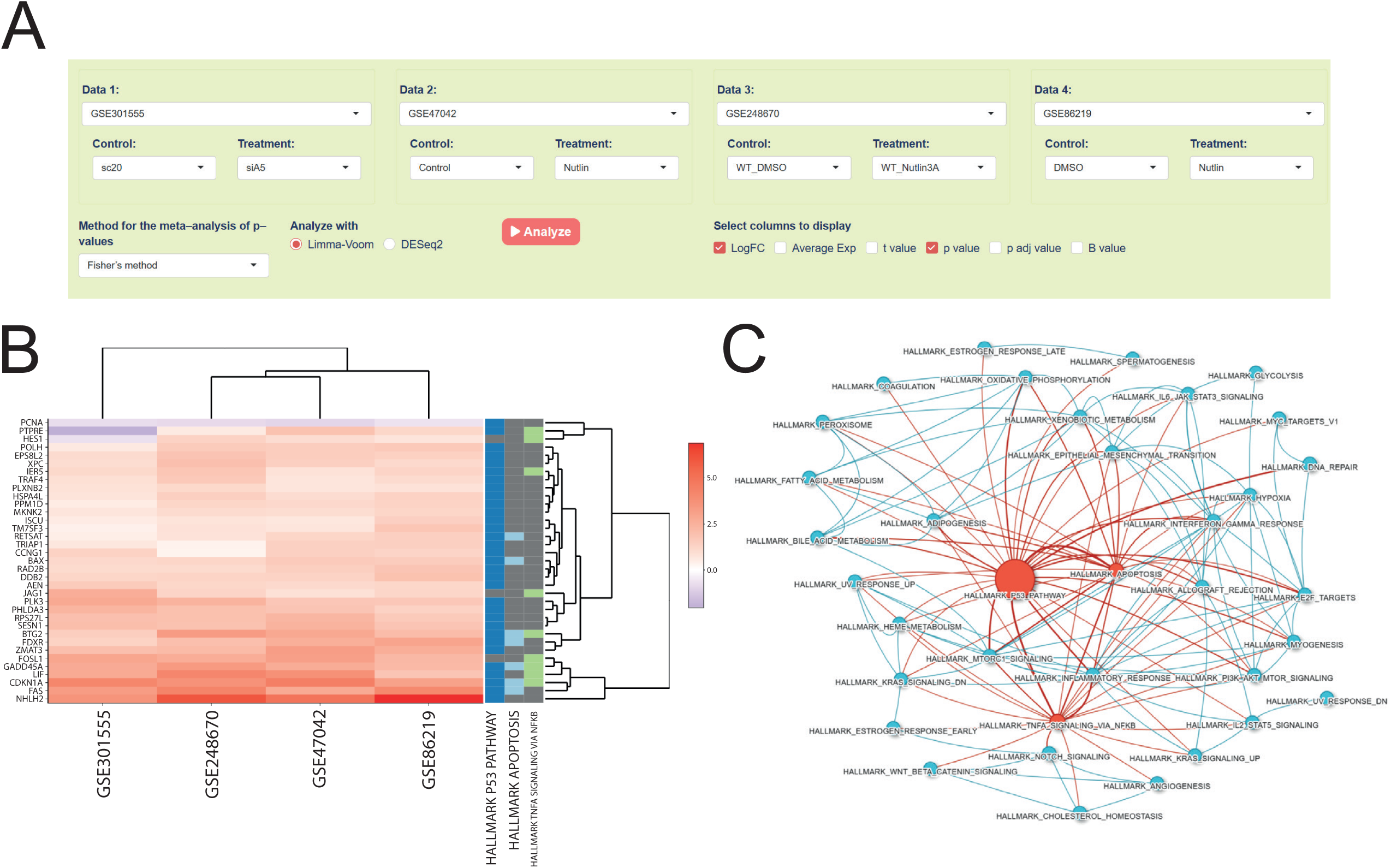
**A)** Selected comparisons and configurations for comparison of siA5 with Nutlin treatments across different datasets. **B)** Heatmap for pathway analysis results on MSigDB Hallmark pathways. **C)** Network created from MSigDB results (q-value < 0.05 for hallmark TP53 pathway, hallmark TNF-A signaling via NFKB, and hallmark apoptosis), where nodes represent pathways and edges represent shared gene numbers between pathways.

According to the GO terms enrichment analysis, enriched terms included cell cycle, apoptosis, and DNA damage repair-related pathways [see Additional File 6], while in TF-gene analysis based on DoRothEA, TP53-related TF-target gene pairs were enriched [see Additional File 7]. Further analysis using the hallmark set of MSigDB resulted in identification of the TP53 pathway, apoptosis, and TNF-α signaling via NF-κB (Figure 3B). To show the capability of TP53-META for better visualization of resulting pathways in MSigDB analysis, a network was generated in which significantly enriched pathways (q-value < 0.05) were highlighted, and their relationships to other pathways were mapped using edge widths reflecting the number of shared genes (Figure 3C).

When comparing treatment clusters by GO-enriched terms, siCHRNA5 treatment uniquely showed enrichment for deaminase activity, distinguishing it from Nutlin treatments. In contrast, terms such as “transmembrane receptor protein tyrosine phosphatase activity,” “transmembrane receptor protein phosphatase activity,” “ubiquitin conjugating enzyme binding,” “damaged DNA binding,” and “small molecule sensor activity” were absent in siCHRNA5 and were not consistently enriched across all three Nutlin treatments either.

Overall our analysis showed that siCHRNA5 treatment was able to induce TP53 targets and genes enriched in commonly known TP53-related processes, demonstrated that siCHRNA5 acted like a TP53 inducer and was in accord with the findings by Cingir Koker et al. (15).

### TP53 independent effects of CHRNA5 depletion in MCF7 cells

We further investigated genes affected by siCHRNA5 in the absence of TP53 by conducting three comparisons:

i. siCHRNA5 vs. control, (ii) siTP53 vs. control, and (iii) siCHRNA5 + siTP53 vs. control. Genes were filtered to include those not significant in the siTP53 vs. control comparison (ii) but significant in the other two (i and iii) (adjusted p < 0.05), yielding 539 genes [see Additional File 8]. Using these genes, we constructed a treatment-gene network with an absolute logFC threshold of 2.5 to further prune and include the highly modulated genes. Our results revealed that *MAP1B*, *DMRTA1*, *CREB5*, and *CDC42EP3* were upregulated in both the siCHRNA5 vs. control and siCHRNA5 + siTP53 vs. control contrasts but showed no association with the siTP53-only node. Conversely, downregulated genes under the same filtering included *CXCR4*, *TMEM270*, *CHRNA5*, *AOX1*, *IGLON5*, *LRG1*, *C1R*, *GLP1R*, and *DLK1* (Figure 4A). We also extracted a C2 gene set from MSigDB (C2: chemical and genetic perturbations) called “Dutertre ESTRADIOL response 24HR UP” (q < 0.05), largely representing secondary, delayed and/or non-genomic Estradiol signaling, and showed that CHRNA5 depletion supported our previous findings, i.e., negative correlation with secondary estrogen signaling (53). Most importantly, this analysis indicated that higher CHRNA5 expression’s positive association with the selected gene set was relatively independent of TP53’s action. Among the TP53-independent siCHRNA5 modulated genes in this set, all were downregulated in the siCHRNA5 vs. control comparison—except for *IL1RAP*. The combined treatment revealed further upregulation of *TPX2*, *NCAPD2*, *TOP2A*, and *SLC29A1* by siCHRNA5 in the absence of TP53 (Figure 4B).

**Fig 4.**
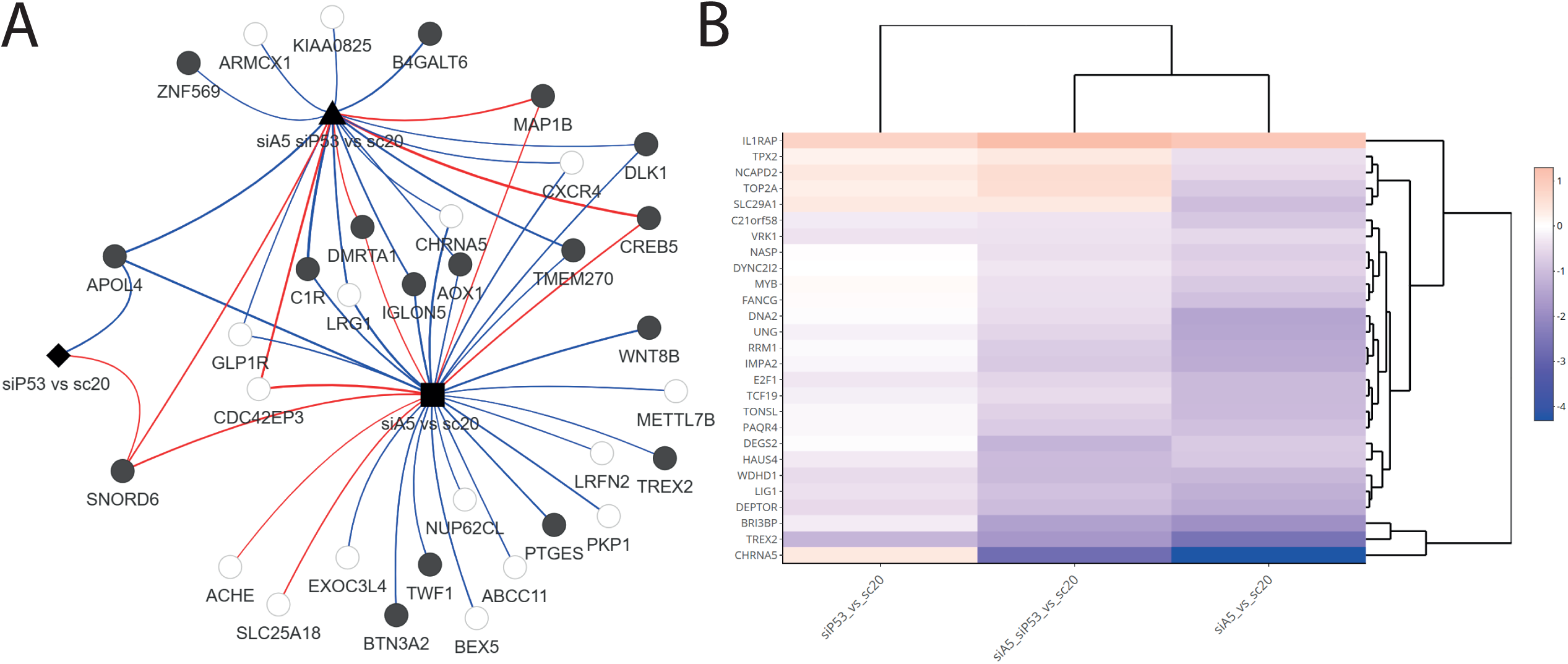
**A)** Treatment-gene network of TP53 independent effects of siCHRNA5 filtered for |logFC| > 2.5. **B)** Heatmap showing logFC values of filtered genes present in MsigDB C2 - “Dutertre ESTRADIOL response 24HR UP.”

### Potential synergistic effect of siCHRNA5 and siTP53 in MCF7 cells

We also identified potential synergistic effects of the combinatorial treatment by comparing four conditions:

i. siCHRNA5 vs. control, (ii) siTP53 vs. control, (iii) siCHRNA5 vs. siTP53, and (iv) siCHRNA5 + siTP53 vs. control. Genes were filtered to include only those significant in the combinatorial treatment (adjusted p < 0.05) but not significant in any of the other comparisons (p ≥ 0.05), to make sure that the combinatorial effect alone was detected stringently. This filtering resulted in 397 genes [see Additional File 9], which were analyzed for functional enrichment using the MSigDB C2: REACTOME gene set. The results revealed enrichment in “ABC family proteins mediated transport” and “Cilium assembly” pathways (p < 0.05, q > 0.05). Among the genes overlapping with the ABC transport pathway, most were upregulated, with the exception of *PSMD10*, *RNF5*, and *ABCB8*. In contrast, genes involved in Cilium assembly were generally downregulated, except for *TMEM216*, *HSP90AA1*, and *TUBA4A* [see Additional File 10].

### Case Study 2: Association between liver fibrosis and TP53

While the tumor suppressor TP53 has been extensively studied in various cancer types, it also plays an active role in non-cancer-related processes, including liver fibrosis. A previous study on mice has shown that TP53 activation in hepatocytes leads to liver fibrosis; on the other hand, lack of MDM2 increases connective tissue growth factor in hepatocytes (54). Further studies have indicated that the early stage of TP53 activation plays an anti-inflammatory and anti-fibrotic role; however, during chronic liver injury, prolonged activation of TP53 promotes differentiation of hepatic stellate cells into the myofibroblasts that produce extracellular matrix (55).

To show the user uploaded external data analysis properties of TP53-META, a meta-analysis of three liver fibrosis datasets, i.e., GSE135251, GSE162694, and GSE174478, was conducted with respect to the TP53 census target gene lists by comparing fibrosis stages 4 and 0 across a total of 125 samples (56, 57, 58). Differential gene expression analysis was performed using Limma-Voom, and meta-p-values were calculated based on the Fisher’s method. Next, genes in the TP53 census target gene list were filtered for a meta p-value < 0.05 threshold and having the same direction in all three datasets. This resulted in a total of 20 TP53-related genes, of which 17 were upregulated in fibrosis stage 4 compared to stage 0, and for three of them (*COBLL1*, *CMBL*, and *MFAP3L*), the expression logFC values were negative (Figure 5A).

**Fig 5.**
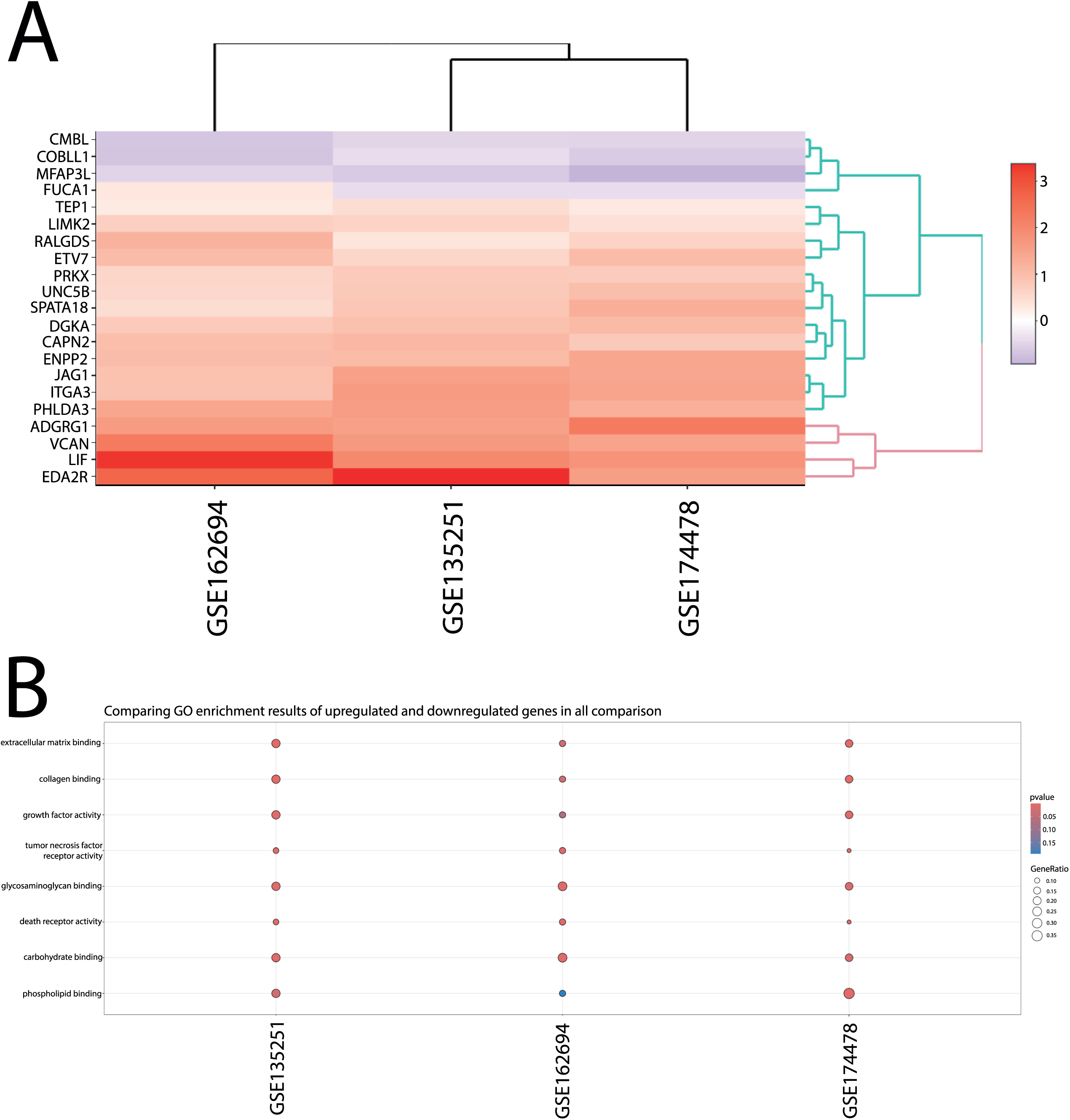
**A)** Heatmap for filtered gene sets in liver fibrosis comparisons with logFC values, where positive values are red and negatives are blue. **B)** GO enrichment results of liver fibrosis comparisons. GeneRatio is depicted as a dot size while the color red indicates smaller p-values.

An interaction between TP53 signaling and other signaling pathways, such as TGF-β and p66Shc, was shown before in the literature (59, 60). To further understand the relationship of resulting TP53 targets in our analysis, we performed enrichment using MSigDB hallmark and C2 gene sets (q value < 0.05). This revealed that “Hallmark Angiogenesis,” “Verrecchia Response to TGFB1 C1,” “IL2 Signaling Up,” and “TAP63 Pathway” were also enriched together with “Hallmark TP53 pathway” [see Additional File 11]. The “Compare Clusters” analysis in the TP53-META further revealed extracellular matrix binding and collagen binding, indicating the upregulation of TP53 targets contributing to the fibrosis process (Figure 5B). Notably, significant differential expression of TP53 between F4 and F0 stages was observed only in GSE162694 (logFC = 0.354), although its downstream targets exhibited consistent expression changes across all three datasets.

While post-hoc filtering for direction concordance is a common practice in transcriptomic meta-analyses, it can sometimes overlook biologically relevant genes that exhibit discordant regulation across datasets (6). Notably, such discordance may be driven by only a subset of studies yet still hold biological significance in general. Several alternative approaches have been proposed to address this challenge (61, 62).

Considering this, we also examined genes in the TP53 target list with significant meta-analysis p-values (meta-p-value < 0.05) without applying directional concordance filters, using the ‘Significant Upregulated’ and ‘Significant Downregulated’ options available in TP53-META. We then utilized the ‘’Treatment–Gene Network’’ feature of the application to visualize genes passing this relaxed filter, i.e., genes with a meta-p-value < 0.05 and | logFC| > 1, resulting in a network of 77 genes (Figure 6).

**Fig 6.**
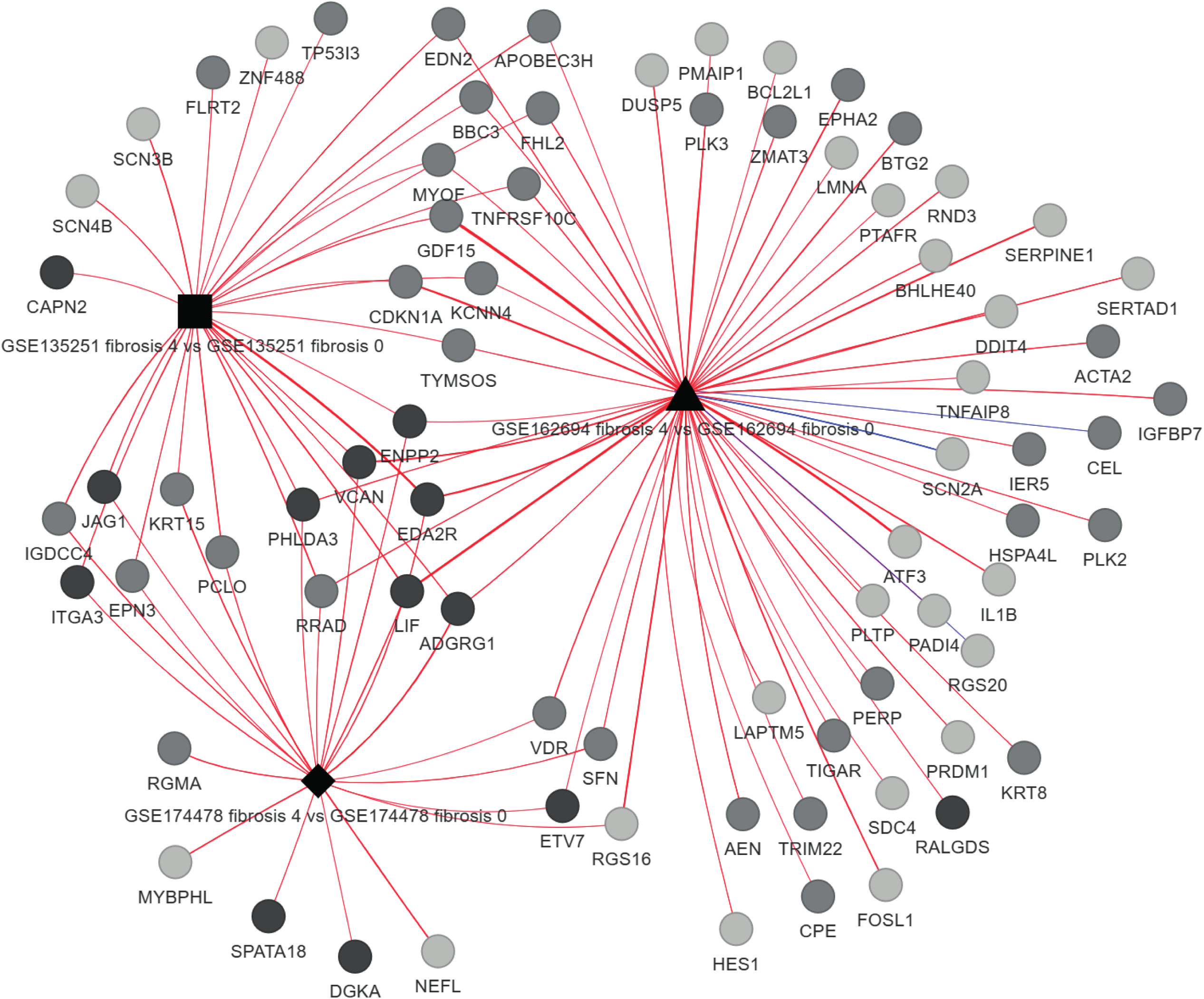
Treatment-gene network of F4 vs F0 comparisons filtered for meta-p-value < 0.05 and |logFC| > 1.

This approach uncovered additional TP53-related genes potentially modulated with the increasing fibrosis stage. Notable examples include *SFN*, *VDR*, and *RGS16*, shared between GSE162694 and GSE174478; *PCLO*, *KRT15*, *EPN3*, and *GDCC4*, shared between GSE135251 and GSE174478; and *MYOF*, *TNFRSF10C*, *TYMSOS*, *KCNN4*, *CDKN1A*, *GDF15*, *FHL2*, *APOBEC3H*, and *EDN2*, shared between GSE162694 and GSE135251.

Considering the role of TP53 in fibrosis, identification of its downstream targets and their role in fibrosis is important for the management of TP53 in the context of liver fibrosis due to its role in anti-fibrosis in the early stages of the disease, while having a turnover for contributing to the disease itself. The gene list provided in this study can be a good candidate list for further understanding the role of downstream TP53 signaling in liver fibrosis, as the literature already points to the importance of TP53 in liver fibrosis.

## DISCUSSION

The current study presents a web-based tool, TP53-META, allowing analysis of TP53-focused datasets through a diverse range of functions across two distinct biological contexts: association between 1) CHRNA5 depletion and induction of TP53 and 2) TP53 target genes and liver fibrosis. Our application stands out among existing tools, as none allow a) visualization of treatment-gene networks and b) comparison of biological themes across the meta-analysis results with individual dataset-level filtering. While tools like Network Analyst (12) and Express Analyst (11) offer strengths in network biology and meta-analysis, they lack the capacity for individual filtering of each dataset, such as identifying genes significant in only one study or applying direction-specific filters within each dataset. They also do not support simultaneous analysis of treatment-gene networks.

Tools such as Omics Playground (9) and OmicsView (10) are more comprehensive in data management and general analysis but lack depth in network-based result visualization. Other meta-analysis tools, such as MAGE and DExMA (63, 64), lack the advanced visualization and network biology features offered by TP53-META. Meanwhile, tools like CAP-RNA-seq, RNA-seqChef, and iDEP support differential gene expression analysis and enrichment with strong visualization capabilities, but they do not support meta-analysis or detailed network creation (65, 66, 67). Moreover, no current tool integrates both TP53-modulated data and curated TP53-related gene lists, thus are not tailored for TP53-specific analyses. A comparison of TP53-META against available tools can be seen in an additional file [see Additional File 12].

Our case studies, uniquely positioned to demonstrate the novelty of TP53-META, enable users to conduct a series of analyses in a TP53-centric context without relying on external tools. In the first case study, we presented a novel in-house RNA-seq dataset with the application, which was analyzed together with publicly available datasets integrated into TP53-META. Our application was able to extract genes consistently regulated by siTP53 treatment or Nutlin across multiple datasets and allowed the visualization and downstream analysis of these gene sets. Furthermore, we demonstrated that CHRNA5 depletion had both TP53-dependent and independent effects, while the simultaneous CHRNA5 and TP53 treatment exhibited synergy. These findings could be important in cancer treatment, in which the TP53 status of patients is known to predict the response to an inhibitor or gene knockdown, as in the integrated dataset from GSE74493 (16). In addition, application of TP53-META to our in-house dataset generated from MCF7 cells treated with siCHRNA5, siTP53, or their combination provided mechanistic evidence for our previous functional studies (15, 53). We also included in TP53-META other combinatorial studies for target gene and TP53 silencing for reanalysis, which can enable users to extract, annotate, and visualize the synergistic or antagonistic effects of combinatorial treatments. Moreover, it is possible for the user to add their own combinatorial study for visualization within a treatment-gene network, which is missing in the literature so far.

In the second case study, the role of TP53 target genes in liver fibrosis was investigated through a meta-analysis where three different studies were integrated, leading to the compilation of results from a total of 125 samples with liver fibrosis stages of F0 or F4. Overall, 20 TP53 targets were significant across three datasets, in which all were upregulated except three. Further analysis of GO terms using the selected genes showed extracellular matrix-related processes, which were in line with the literature (68, 55). While no study has specifically focused on TP53 target gene expression across fibrosis stages, there has been a continuing effort to identify genes associated with liver fibrosis. One such example is steatoSITE, developed by Kendall et al. (69), which allows users to explore transcriptomic changes in liver fibrosis. Using this tool, we examined the direction of change in our resulting gene list from TP53-META and found that our results were consistent with those in steatoSITE.

Overall, TP53-META was successful in comparing different datasets in TP53-related contexts, where it was able to integrate existing or external datasets for differential expression analysis, functional profiling, use of DoRothEA and MSigDB sets, as well as cluster comparison. It also allowed the creation of treatment-gene networks, calculation of correlations, computation of meta-p-values using six different methods, and application of directional filtering. Despite these capabilities, it is important to acknowledge that TP53-META still has several limitations. Future versions should incorporate user-defined preprocessing options, provide greater control over the computation of meta-p-values, and expand the collection of TP53-modulated datasets, gene sets, and analytical functions. Enhancements in performance metrics and overall scalability of the application would also further strengthen its utility.

## CONCLUSIONS

TP53-META offers a platform to compare and visualize experiments within or between datasets with the calculation of meta-p-values and filtering for directionality. Our web tool filled the gap for a comprehensive and modular RNA-seq analysis tool focusing on TP53 by enabling differential gene expression analysis, functional analysis, and visualization of gene-treatment interactions of integrated public or in-house datasets simultaneously. The upload function further demonstrated the unique capability of TP53-META to meta-analyze user uploaded datasets within the context of TP53 target gene sets. TP53-META is available at <u>konulabapps.bilkent.edu.tr:3838/TP53-Meta1.5/</u>

## AVAILABILITY AND REQUIREMENTS

**Project Name:** TP53-META

**Project Home Page:** http://konulabapps.bilkent.edu.tr:3838/TP53-Meta1.5/

**Operating System:** Platform independent.

**Programming language:** R.

**Any restrictions to use by non-academics:** None.

## LIST OF ABBREVIATIONS

TF: Transcription factor
DGE: Differential gene expression
MsigDB: Molecular signatures database
PCA: Principal component analysis
GSEA: Gene set enrichment analysis
GO: Gene ontology
FDR: False discovery rate
HSCs: Hepatic stellate cells
ECM: Extracellular matrix
MFBs: Myofibroblasts
DGEA: Differential gene expression analysis

## DECLARATIONS

### Ethics approval and consent to participate

Not applicable.

### Consent for publication

Not applicable.

### Availability of data and materials

All datasets used in TP53-META are available online at the GEO (Gene Expression Omnibus) database (https://www.ncbi.nlm.nih.gov/geo/). GSE301555 (https://www.ncbi.nlm.nih.gov/geo/query/acc.cgi?&acc=GSE301555) GSE47042 (https://www.ncbi.nlm.nih.gov/geo/query/acc.cgi?acc=GSE47042) GSE173364 (https://www.ncbi.nlm.nih.gov/geo/query/acc.cgi?acc=GSE173364) GSE68248 (https://www.ncbi.nlm.nih.gov/geo/query/acc.cgi?acc=GSE68248) GSE139546 (https://www.ncbi.nlm.nih.gov/geo/query/acc.cgi?acc=GSE139546) GSE83635 (https://www.ncbi.nlm.nih.gov/geo/query/acc.cgi?acc=GSE83635) GSE84877 (https://www.ncbi.nlm.nih.gov/geo/query/acc.cgi?acc=GSE84877) GSE74493 (https://www.ncbi.nlm.nih.gov/geo/query/acc.cgi?acc=GSE74493) GSE124508

(https://www.ncbi.nlm.nih.gov/geo/query/acc.cgi?acc=GSE124508) GSE59841 (https://www.ncbi.nlm.nih.gov/geo/query/acc.cgi?acc=GSE59841) GSE86219 (https://www.ncbi.nlm.nih.gov/geo/query/acc.cgi?acc=GSE86219) GSE248670 (https://www.ncbi.nlm.nih.gov/geo/query/acc.cgi?acc=GSE248670) GSE89110 (https://www.ncbi.nlm.nih.gov/geo/query/acc.cgi?acc=GSE89110)

## Competing interests

The authors declare that they have no competing interests.

## Funding

This project received funding from TUBITAK with grant number 219Z029 and from the European Horizon’s research and innovation program HORIZON-HLTH-2022-STAYHLTH-02 under agreement No 101095679. Funded by the European Union. Views and opinions expressed are however those of the author(s) only and do not necessarily reflect those of the European Union. Neither the European Union nor the granting authority can be held responsible for them.

## Authors’ contributions

OK conceived and supervised the study. AGK and ST performed the wetlab experiments and AGK and AMA conducted data-analyses related to TP53 and CHRNA5 siRNA treatments of MCF7 cells. MVO initially developed the TP53-META application; and EK, AMA, TD and OK further optimized and enhanced the web application; EK performed data analyses of the case studies and obtained the figures. EK, AMA, OK wrote and all authors read, revised and approved the manuscript.

## ADDITIONAL FILES

| File Name | File Format | Title of data | Description of data |
| --- | --- | --- | --- |
| Additional File 1 | PDF | qPCR and western blot results of siRNA treatments. | Validations of siCHRNA5, siTP53, and combination treatments in MCF7 cells using (A) qPCR and (B) western blot. |
| Additional File 2 | PDF | Common genes of TP53 depletion and TF analysis | (A) Venn diagram of genes significant in each study and (B) heatmap of DoRothEA results using the resulting 94 genes. |
| Additional File 3 | PDF | DisGeNET analysis of TP53 depletion | Heatmap of DisGeNET results obtained by using logFC values of |
|  |  |  | GSE301555 |
| Additional File 4 | PDF | GO terms enrichment analysis of TP53 depletion | Enrichment map of GO terms enrichment analysis on logFC values of GSE301555 |
| Additional File 5 | PDF | Correlation between siCHRNA5 and Nutlin treatments | Correlation matrix showing Pearson correlation between Nutlin-treated datasets and siCHRNA5 treatment |
| Additional File 6 | XLSX | Go terms enrichment analysis of siCHRNA5 | A table containing the results of the GO terms enrichment analysis on 94 genes obtained from siCHRNA5 samples and Nutlin-treated datasets |
| Additional File 7 | XLSX | DoRothEA results | A table containing |
|  |  | of siCHRNA5 | the results of DoRothEA on 94 genes obtained from siCHRNA5 samples and Nutlin-treated datasets |
| Additional File 8 | XLSX | Independent effect gene list | DGEA results of filtered genes for TP53 independent effects of siCHRNA5, including adjusted p-values and logFCs |
| Additional File 9 | XLSX | Synergistic effect gene list | DGEA results of filtered genes for synergistic effects of siCHRNA5 and siTP53, including adjusted p-values and logFCs |
| Additional File 10 | PDF | Enriched terms of the synergistic | Heatmap showing logFC values of |
|  |  | gene list | genes enriched in<br>MsigDB C2:<br>REACTOME gene<br>set for synergistic<br>gene list. The<br>enriched terms<br>include “ABC family<br>proteins mediated<br>transport” and<br>“Cilium assembly” |
| Additional File 11 | XLSX | Enrichment of<br>fibrosis-related<br>TP53 targets in<br>MSigDB | A table containing<br>enriched terms in<br>MSigDB H and C2<br>gene sets for<br>fibrosis-related<br>TP53 targets |
| Additional File 12 | XLSX | Comparison of<br>TP53-META with<br>other applications | A table containing<br>comparisons of<br>TP53-META<br>against 7<br>applications under<br>7 broad categories. |

## Notes

https://www.ncbi.nlm.nih.gov/geo/query/acc.cgi?acc=GSE301555

### Competing Interest Statement

The authors have declared no competing interest.

