## Supplementary figures and images for "TP53-META, a meta-analysis tool for comparative transcriptomics of TP53 dependency: Examples from target silencing and liver fibrosis"

### Additional File 1

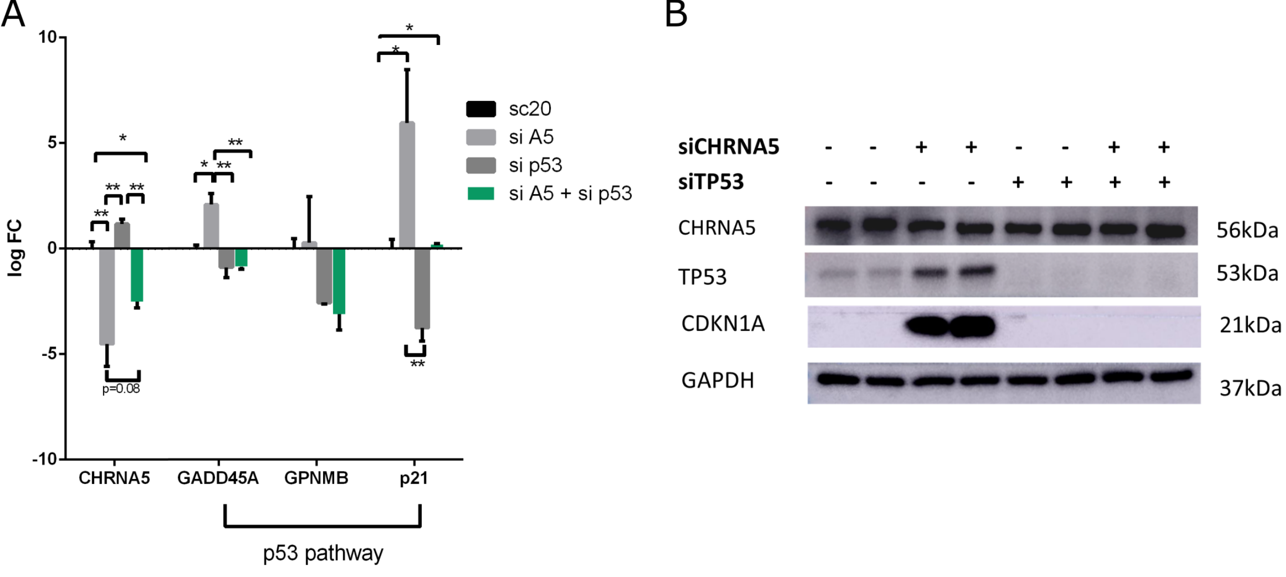

### Additional File 2

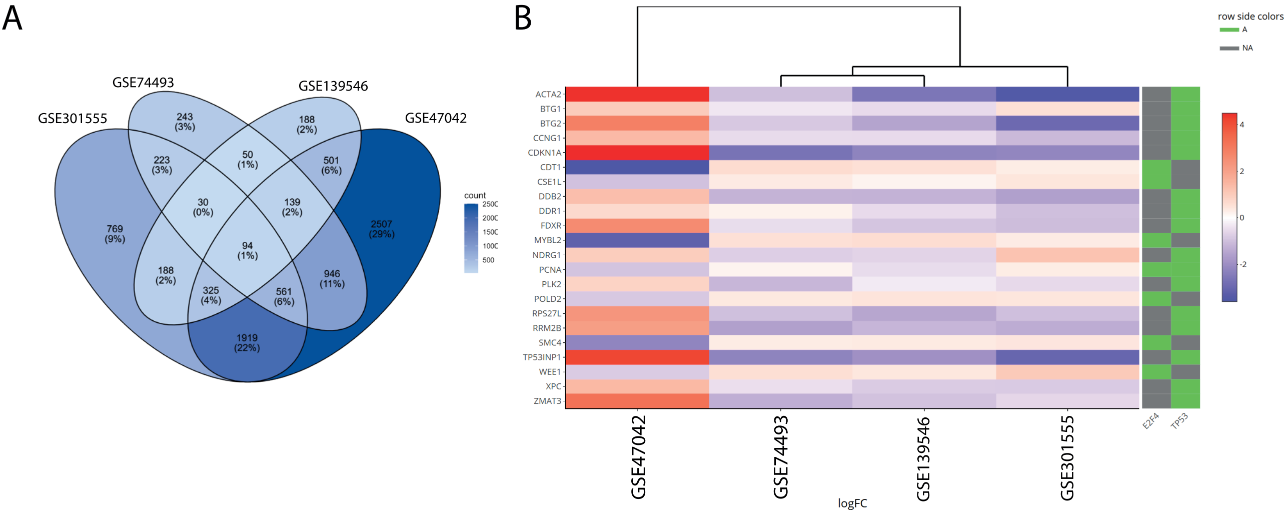

### Additional File 3

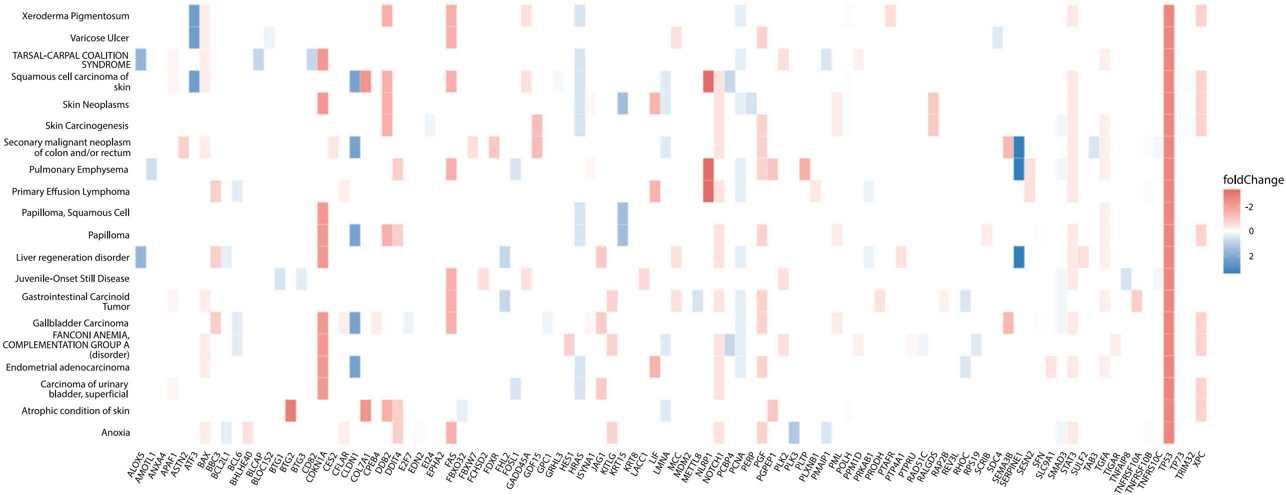

### Additional File 4

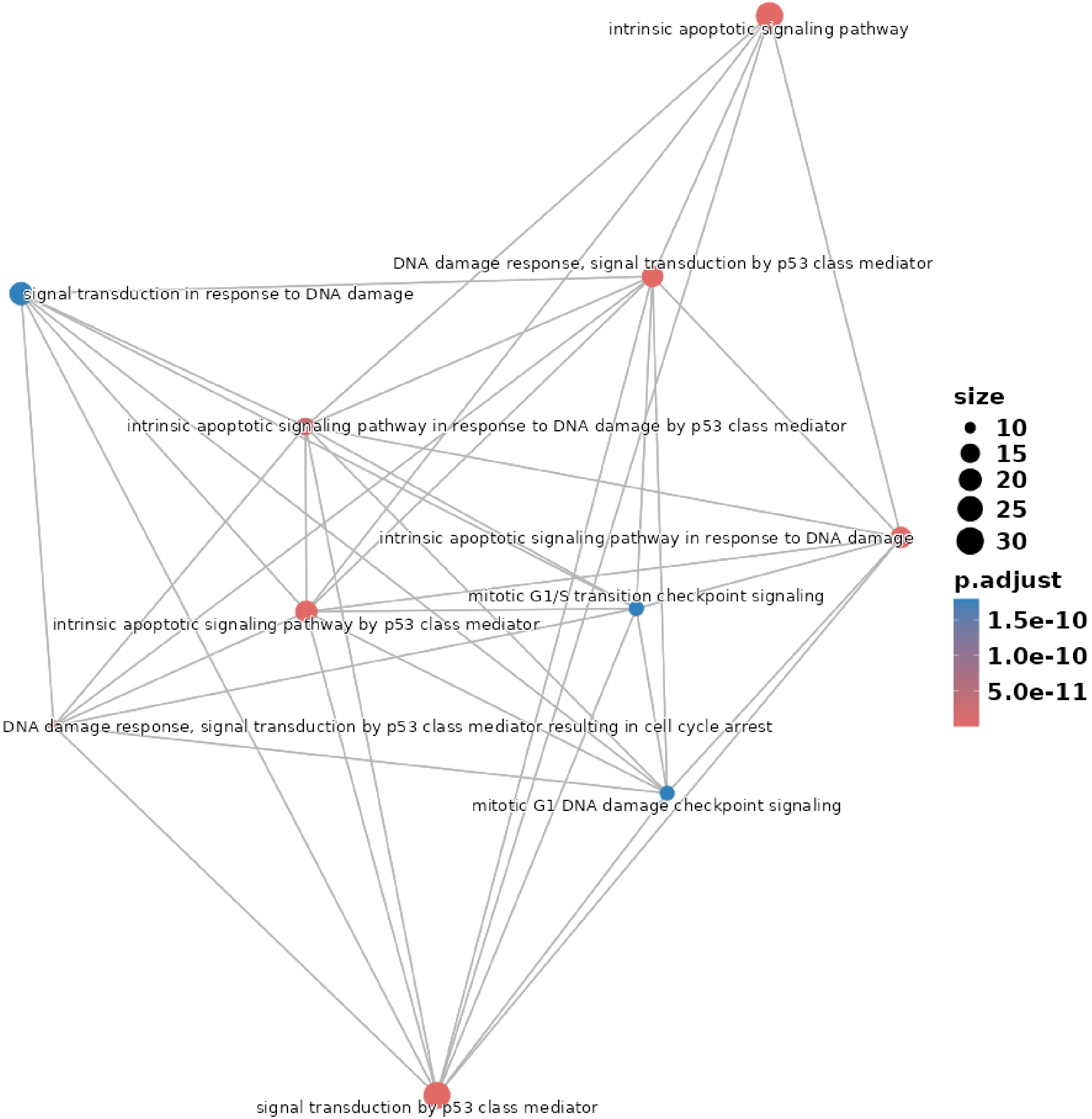

### Additional File 5

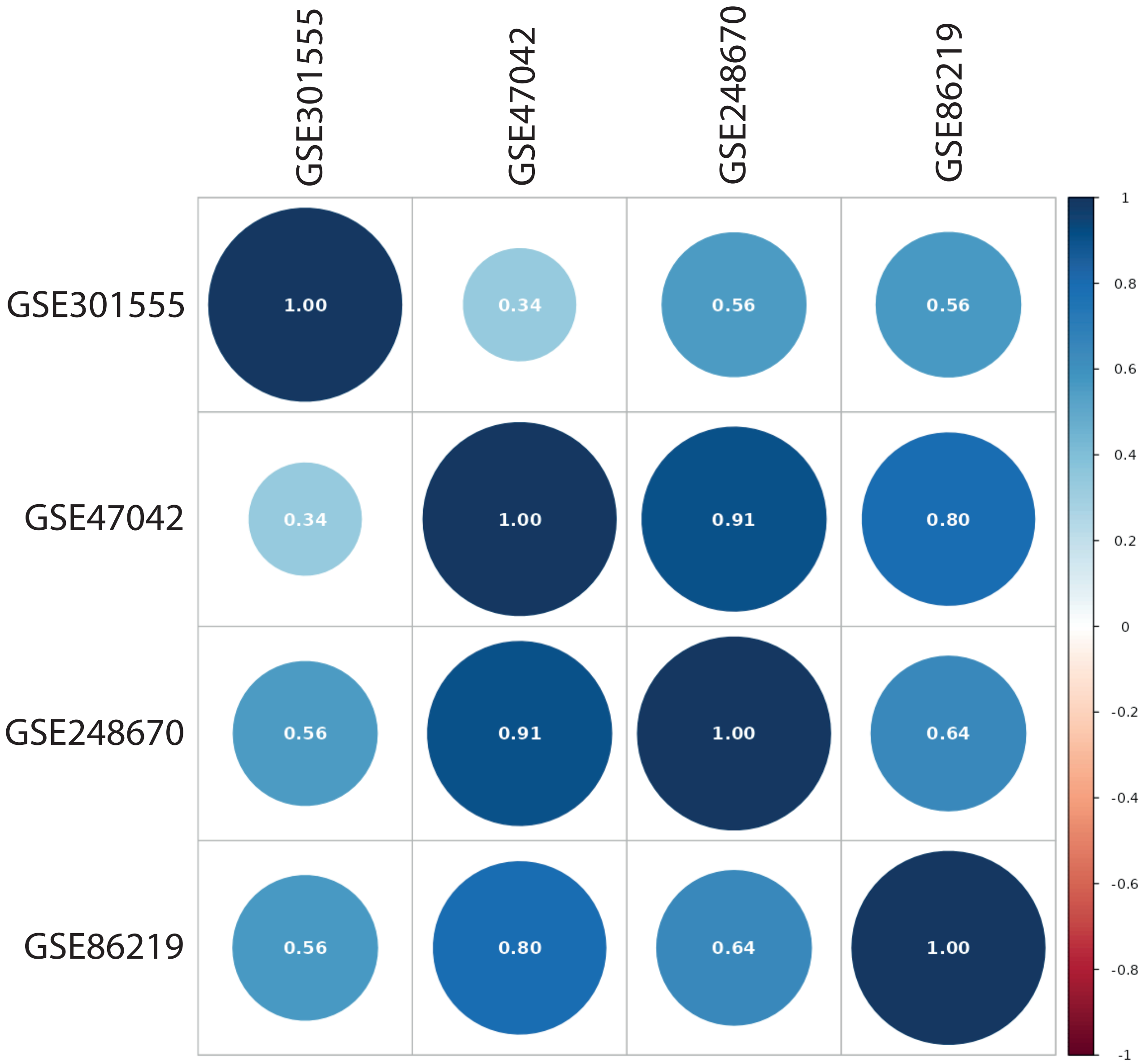

### Additional File 10

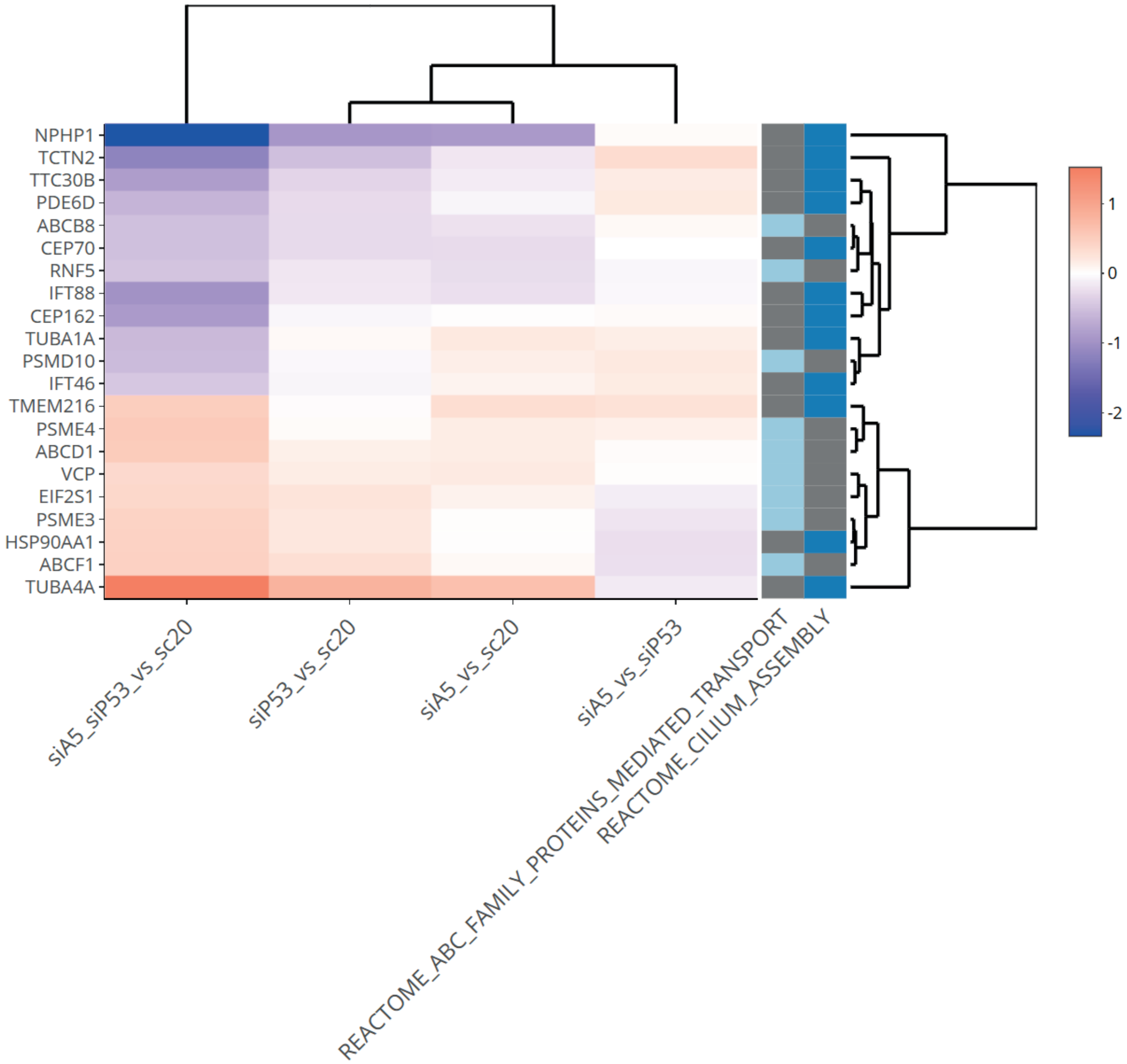
